# Environmental Correlates for Anticoagulant Resistance in house mice *Mus musculus*

**DOI:** 10.64898/2026.04.12.718056

**Authors:** Luigi F. Richardson, Mehra Balsara, Celine Larose, Catherine I. Cullingham, Albrecht I. Schulte-Hostedde

## Abstract

Genetic resistance to anticoagulant rodenticides is a world-wide evolutionary phenomenon among invasive commensal rodents, that has not been investigated in Canada. We sequenced exon 3 of the VKORC1 gene of house mouse *Mus musculus* samples obtained from pest management professionals in Ontario and Quebec, to search for mutations known to confer resistance. Sanger sequencing was used alongside a novel qPCR assay for codons 128 and 139. We detected high prevalence (99%) of two non-synonymous single nucleotide polymorphisms in house mice, previously found in the USA and Europe, known to cause extreme resistance to first and some second-generation anticoagulants (L128S & Y139C). Homozygous resistant mice were more common in high population density areas. L128S alleles were more common in Southwestern Ontario, and Y139C in Central Ontario, despite high linkage disequilibrium. Detection success was far greater with qPCR than with Sanger sequencing. We conclude that uncoordinated rodenticide usage has selected for extreme resistance in mice throughout Ontario. Therefore, chemical control of the house mouse may be ineffective with first-generation anticoagulants throughout Ontario. This suggests that the evolution of wild urban mice is influenced by pest management practices, which may vary by region within the province.

## Introduction

Invasive pests are responsible for significant ecological, social and economic losses worldwide (Pimentel et al. 2000), particularly in urban ecosystems, where they have caused almost $330 billion USD in damage within the past 60 years (Cuthbert et al. 2022; Heringer et al. 2024). Commensal rodents are seldom found far from human settlements and rely on human food sources and infrastructure (Doherty et al. 2016), and are thus quintessential invasive species (Weissbrod et al. 2017). Commensal rodents include the black rat *Rattus rattus*, Norway rat *Rattus norvegicus* and the house mouse *Mus musculus*. These species frequently out-compete native and ecologically similar species in human-altered environments (White et al. 2026), and may be linked to nearly 45% of recent terrestrial vertebrate species extinctions (Gorbunova et al. 2008; Vinyard and Payseur 2008). House mice are likely the world’s most widespread invasive commensal, inhabiting every continent except Antarctica, including islands of the sub-Antarctic region (Renaud et al. 2013). They are seldom studied in the wild (Boell and Tautz 2011), but cause significant economic damage (Pimentel et al. 2000) and carry numerous pathogens (Meerburg et al. 2009). For these reasons, efficient control methods are needed.

Anticoagulant rodenticides (ARs) were successfully used for house mouse control worldwide in the earlier half of the 20^th^ century (Campbell and Link 1941). As a result of intensive usage, resistance to first-generation ARs (FGARs) in target species, including house mice, has evolved in numerous regions, rendering them ineffective for control (Pelz et al. 2005). Anticoagulants are structural analogs of vitamin K, effectively blocking vitamin K from entering the vitamin K epoxide reductase receptor (VKORC1) and preventing clotting in vertebrates (Smith et al. 1993). Resistance in commensal rodent populations is largely the result of natural selection for single-nucleotide polymorphism (SNP) mutations that alter the amino acid composition of VKORC1 preventing ARs from binding with vitamin K receptors, but still allow vitamin K binding, needed for the creation of clotting factors (Rost et al. 2004). For example, a A/G SNP in codon 139 changes Tyrosine to Cysteine. Individuals homozygous for the Cysteine mutation increase the LD50 for bromadiolone, a popular second-generation AR (SGAR), by more than ten times (Baxter et al. 2022). Heterozygotes express less resistance than homozygotes, and increased life expectancy relative to homozygotes which often suffer from vitamin K deficiency (Baxter et al. 2022).

Although allelic and genotypic diversity for VKORC1 resistance SNPs often differs by geographic region (Díaz and Kohn 2021), few authors have examined the relationship between the prevalence of resistance SNPs and different ecological and landscape factors. For example, the prevalence of AR resistant SNPs should be associated with socioeconomic and land use factors that predict high AR use because of natural selection (Feng and Himsworth 2014; Hindmarch et al. 2018). Scientists may be able to determine how AR resistance in house mouse populations is evolving in response to rodenticides (Haniza et al. 2015), and associate this with factors that are correlated with AR usage (Tripodi et al. 2024).

Since urbanized regions (Haniza et al. 2015) and those with lower average income (Tripodi et al. 2024) may experience more house mouse infestations, and use AR chemicals more often for control, this may create an urban-rural socioeconomic gradient of rodenticide resistance. Alternatively, regions with lower income may have less access to rodenticides, which could make resistance less prevalent. Commensals are severely understudied, and research on resistance can further our understanding of how pest management as a selective pressure against invasive species in novel anthropogenic ecosystems (Murray et al. 2025).

Our understanding of AR resistance in a Canadian context is limited to no-choice feeding tests on laboratory-held wild mice conducted in the 1980s (Siddiqi and Blaine 1982). Here, we survey house mice southern Ontario and western Quebec for the presence of known SNPs/alleles associated with AR resistance (Pelz et al. 2005). We also use socioeconomic and land use data to test the prediction that areas with presumed elevated use of ARs will have greater prevalence of the AR resistance alleles. Finally, we used Sanger sequencing and a novel qPCR assay for all samples to compare detection success between both methods.

## Methods

### Field Methods

Tissue samples from 79 house mice were collected by pest management companies from 16 locations around Ontario (and one in Gatineau, Quebec) between February and October 2024. Samples were obtained from commercial or residential clients and stored in domestic freezers at the company’s office for less than three months before pickup and processing. We were provided with the exact address, closest main intersection or postal code to indicate location of capture for all samples donated by the pest management companies. Upon collection from pest management companies, all specimens were stored at -20°C at Laurentian University.

### Independent variables

Census tract data (CensusMapper: OpenStreetMap, https://censusmapper.ca/) were used to determine socioeconomic and demographic metrics including population density, median income, and education rates for the areas in which each collection site was located, for all 16 collection sites. Census tracts in Ontario are good proxies for neighbourhoods as a result of their size and social uniformity (Pham 2024).

The respective area of four land types encapsulating capture sites were recorded using the ArcGIS® online polygon tool: industrial or commercial (factories, stores, malls, schools, churches, apartment buildings, etc.), residential (detached, semi-detached or row/town homes), agricultural (cultivated fields) and natural (wild/greenspaces, ArcGIS Online 2025). All lands surrounding the capture site within the confines of four-lane roads or other “impassible” barriers (Rico et al. 2007; McGregor et al. 2008) were assessed. If only postal code was provided, all lands within the area of the postal code were assessed. The percentage of developed land (residential + commercial + industrial) was then used as a proxy for rodenticide usage intensity, and to predict the presence/absence and percentage of resistant rodents. We hypothesized that rodenticide usage intensity would be greater in more “urbanized”, densely-populated and lower-income areas (Hindmarch et al. 2018; Murray et al. 2025), as would resistance. However, we also considered that an inverse relationship could exist, in that some urbanized areas may be lower in average income (Pham 2024), which may preclude widespread rodenticide usage, and hence, selection for resistance alleles.

### Laboratory Methods

A small sample (approx. 1cm) of tail or ear tissue was collected from the carcasses and frozen at -20°C, or stored in ethanol for later DNA extraction and PCR amplification of the VKORC1 gene (Rost et al. 2009). The standard DNeasy Blood and Tissue Kit (Qiagen, Mississauga) was used for DNA extraction following the manufacturers protocol. Exon 3 of the VKORC1 was amplified using the forward primer from (Ruiz-López et al. 2022): TTTCACCAGAAGCACCTGCTGYC. Sequencing occurred in two rounds. The first round was only run for five samples to test the quality of the sequences. PCR conditions for this round of sequencing included 1 U of standard Taq Polymerase, 1x buffer, 0.6 mM dNTP, 0.2 uM of forward and reverse primers, and 10 ng DNA in a 20 uL reaction. Cycling conditions were run as follows: 95°C for 2 mins, 30 cycles of 95°C for 30 seconds, 56 °C for 1 min, 72 °C for 1:30 min, followed by 72 °C for 5 min, and held at 10 °C. The remaining samples were amplified during the second round of sequencing. PCR conditions were consistent with the first round with the following exceptions: 1 U of Platinum™ Taq Green Hot Start DNA Polymerase was used instead of standard Taq Polymerase and 20 ng DNA used in a 15 ul reaction. For the second round the cycling conditions were 94°C for 2 mins, 30 cycles of 94°C for 30 seconds, 56 °C for 30 sec, 72°C for 1 min, followed by 72 °C for 2 min, and held at 10 °C.

PCR products were then analyzed for fragment size, and overall quality, using agarose gel electrophoresis. Samples were treated with ExoSAP-IT (Applied Biosystems, Mississauga) to remove free dinucleotides and remaining forward primer was added before being shipped to The Center for Applied Genomics (SickKids, Toronto) for Sanger sequencing. The resulting chromatograms were trimmed and checked for quality by manual inspection using Finch TV v. 1.5 (Geospiza 2006). Alignments were then performed with the *M. musculus* (NC_000073.7) “wild-type” in UGENE v.52.0. The presence of resistance alleles was also completed manually by searching for SNPs in comparison with the wildtype sequence, with a specific focus on codons 128 and 139 (hotspots for resistance SNPs, (Rost et al. 2009).

### Quantitative PCR assay

Following screening of the successfully sequenced individuals, we developed a qPCR assay to target the resistance SNPs identified. Primers specific to the *Mus musculus* VKORC1 gene (NC_000073.7), along with allele-discriminating probes for each SNP, were designed and evaluated using PrimerQuest^TM^ and OligoAnalyzer ^TM^ (IDT, Coralville, Iowa, USA https://www.idtdna.com/SciTools.), as well as Primer-BLAST (https://www.ncbi.nlm.nih.gov/tools/primer-blast/) to minimize dimer formation and optimize assay conditions (Tab. 1). Quantitative PCR reactions were plated in 96-well plates, consisting of 5 μL of DNA (ultrapure H_2_O was used for controls) and 15 μL of reagent cocktail (Tab. 2). All house mouse samples were plated in duplicate for each assay. Plates were sealed with an adhesive film and centrifuged (Eppendorf bucket centrifuge 5810R 15 amp version) for 2 minutes at 880 rpm/rcf at 19 □C. Amplification was performed using the QuantStudio 3 Real-Time PCR Instrument (96-Well 0.2 ml Block) using the manufacturers conditions (Applied Biosystems TaqPath ProAmp Master Mix, Applied Biosystems 2026) as follows: a pre-read at 60 □C for 30 seconds, initial denature/enzyme activation at 95 □C for 5 minutes, followed by 40 cycles of denaturation at 95 °C for 15 seconds and annealing/extension at 60 □C for 60 seconds, and finally a post-read at 60 □C for 30 seconds. After completion, the data was analyzed using Diomni^TM^ Design and Analysis (RUO) 3.1.0 software (Applied Biosystems 2026, https://www.thermofisher.com/ca/en/home/life-science/pcr/real-time-pcr/diomni-design-analysis-software.html). The amplification plot was used to check quality, while the allelic frequency plots showed distinct clustering of genotypes (Tab. 1). Composition of the PCR reagent mixtures are given in Table 2.

**Table 1:**
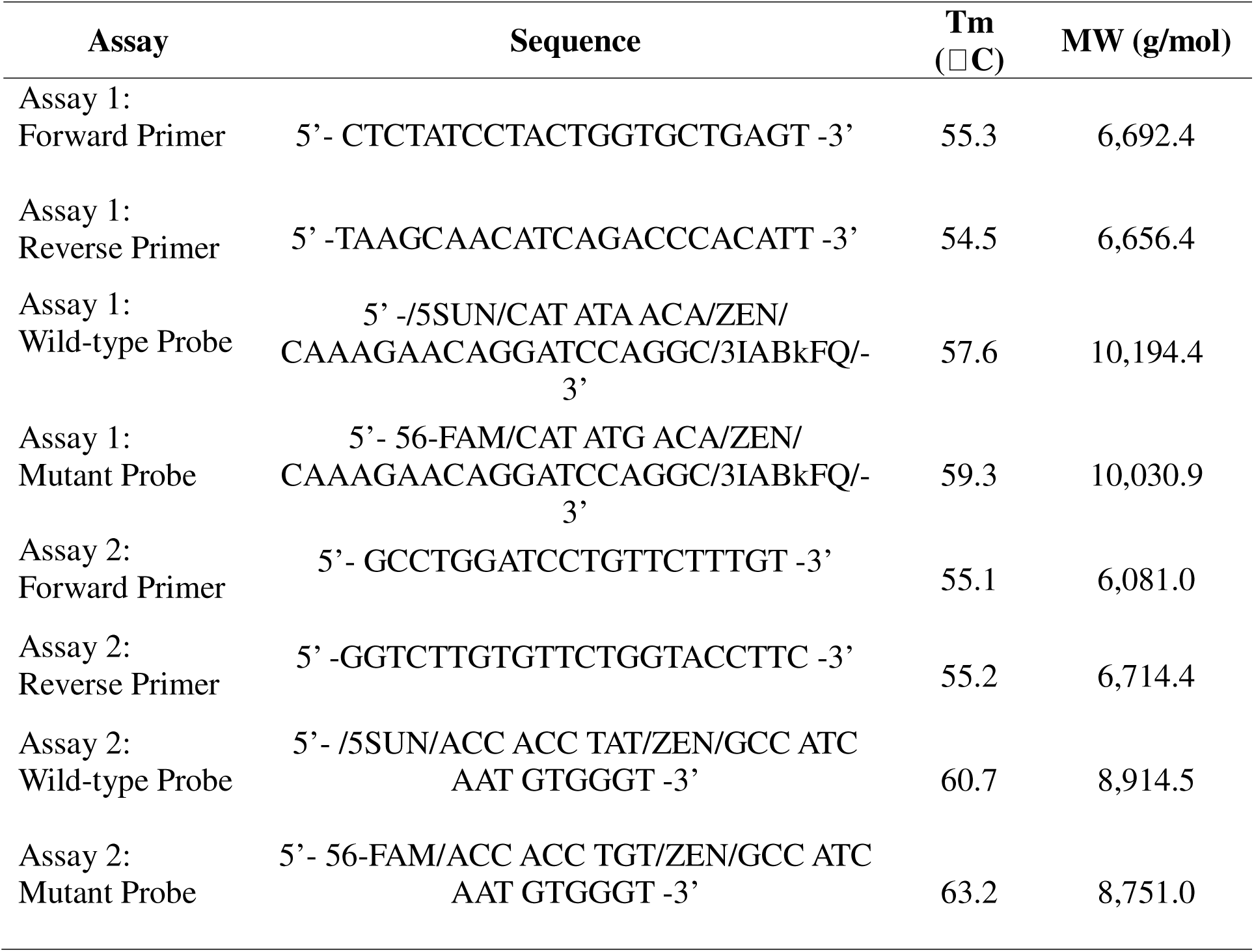
Primer and probe sequences, melting temperatures (Tm), and molecular weights (MW) for TaqMan qPCR assays targeting VKORC1 L128S (Assay 1) and Y139C (Assay 2) SNPs

**Table 2:** Composition of the PCR reagent mixture for 15 μL qPCR reactions

| Reagent | Volume/Reaction |
| --- | --- |
| 4 X Master Mix | 5 µL |
| 10 µM Forward Primer | 1 µL |
| 10 µM Reverse Primer | 1 µL |
| 10 µM Wild-Type Probe | 0.5 µL |
| 10 µM Mutant Probe | 0.5 µL |
| Ultrapure H2O | 7 µL |

### Statistical Methods

Statistical analyses were performed in R Studio (R Core Team 2024, R version 4.4.2). Analyses, models and figures were created with packages “genetics”, “HardyWeinberg”, and “ggplot2”.

We used contingency tables with Fisher’s tests to determine if different regions of Southwestern Ontario, Central Ontario and Quebec varied in terms of resistant and wildtype allele and genotypic frequencies. We tested for Hardy-Weinberg equilibrium and compared observed to expected genotypic frequencies using Haldane’s exact test to determine if resistant loci had unexpected genotypic frequencies within regions. Finally, we calculated D’ for Pairwise linkage disequilibrium and used Chi-square to determine the frequency and significance of resistant allele co-occurrences between different regions.

Correlates used to model resistance linearly were post-secondary education rate (% of census tract population), population density (people/km^2^), median annual household income ($CAD/year), and the percentage of developed land within connected habitat at the capture site (ArcGIS Online 2025; CensusMapper 2025). To assess the relationship between the variables that served as proxies for AR use, we used log-log linear regressions with Gaussian distributions. We then combined these correlates with PCA to capture covariation as a singular linear predictor variable. PC1 was used to predict the percentage of homozygous resistant individuals at a capture site. This method was used in addition to single linear models of each predictor partially due to multicollinearity caused by multiple regression. It also allowed us to determine the effects of all predictors combined and easily interpret their covariation in accordance with the response variable. All model residuals were checked for normal distributions with Shapiro-Wilk tests, for homoskedasticity with Breusch-Pagan tests, and Cook’s distance was used to locate any excessively influential data points.

## Ethics Statement

All commensal rodent samples were collected by certified pest management professionals in the employ of licensed extermination companies in the province of Ontario, as a part of their regular employment duties and with the permission of management. Sex and age of rodent samples was not recorded.

Open AI’s ChatGPT was used to assist in writing R studio scripts for data analysis, which were subsequently checked for correctness and modified for customization.

## Results

A total of 79 house mice were donated from pest management companies between February and October 2024. A total of 52 of these samples (GenBank acc. no. PX092197-PX092307) produced readable chromatograms for exon 3 from Sanger sequencing of the VKORC1 (66% success). Readable chromatograms were produced for the region between Codons 120 through 142. This 66-base-pair section of exon 3 contained no segregating sites other than 128 and 139 (Tab. 3). These samples were amplified using both qPCR assays (128 and 139), showing complete concordance with the chromatogram results. An additional 27 samples which did not produce readable chromatograms, were successfully amplified using the developed qPCR assays allowing for genotype detection at codons 128 and 139 in exon 3 (100% qPCR success). Samples were from 16 distinct locations around Ontario and Quebec, including Northern Ontario (two individuals, one site), Central Southern Ontario (52 individuals, 12 sites), Southwestern Ontario (18 individuals, two sites), and Quebec (or “Eastern”, seven individuals, one site in Gatineau, near Eastern Ontario Border, Fig. 1).

**Figure 1:**
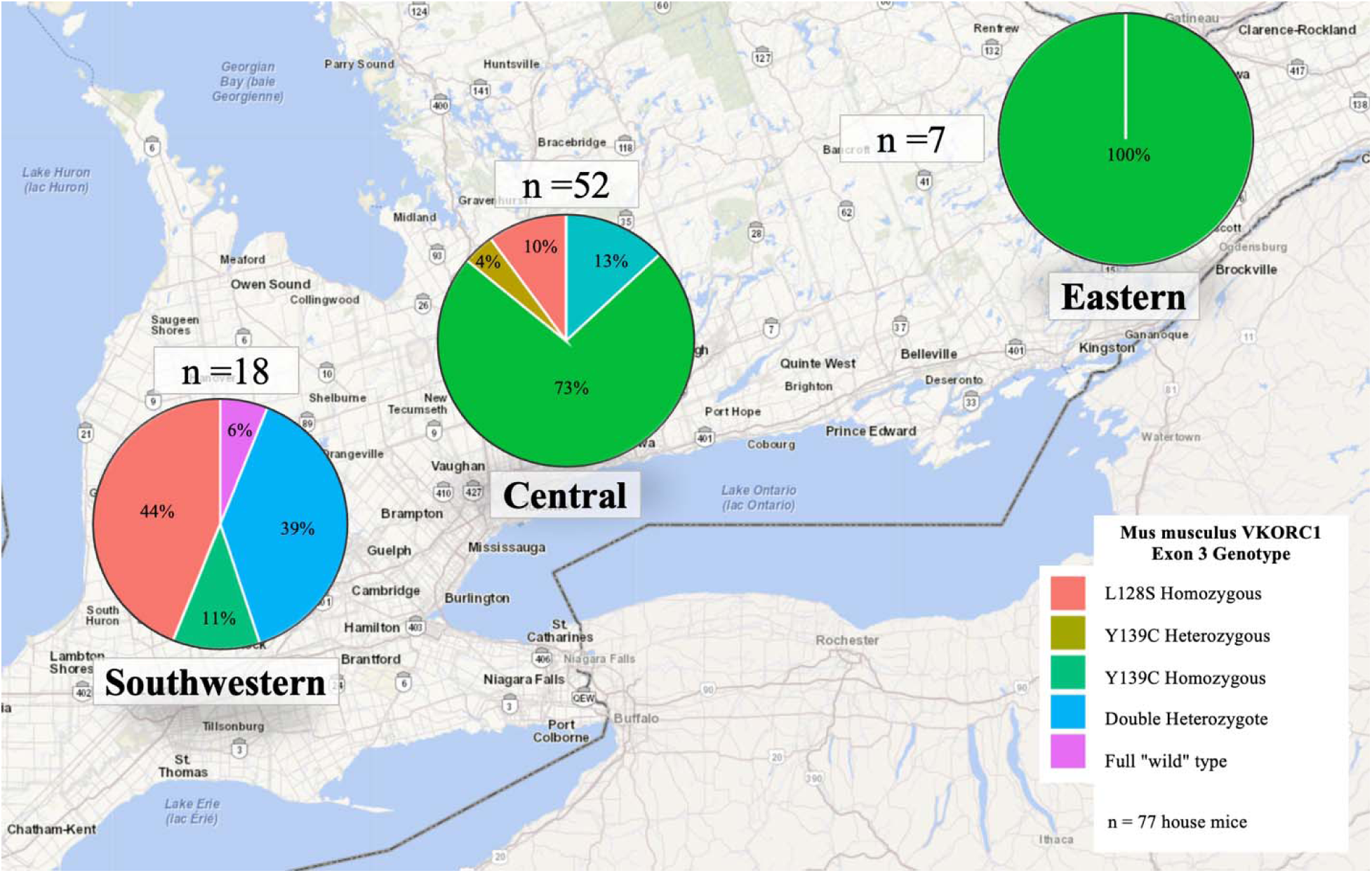
Genotypic frequency comparison between regions of Ontario. Central Ontario and Eastern (Gatineau) had significantly more Y139C homozygotes than Southwestern. Sudbury (Northern Ontario), which had an additional two samples, is not shown here, hence the sample size of 77.

**Table 3:** Allelic counts and frequencies across regions. Allele frequencies differed between all regions (except between Central and Eastern). Fisher’s p column contains values for Fisher’s test of all allele frequencies between the regions being compared (i.e. SW vs C). Bonferroni comparisons are denoted by grey shading and asterisks: L128S was more common in Southwestern, Y139C in Central. % Total denotes the number of individuals with a copy of only one of the two mutations, including Northern Ontario (two additional samples).

|  | **S. Western** | **Central** | **Eastern** | **TOTAL** | **Fisher's p** |  |
| --- | --- | --- | --- | --- | --- | --- |
| Tyr139Cys | 15% (11) | 41% (85)* | 50% (14) | % (111) | *SW vs C* | < 0.001* |
| Leu128Ser | 32% (23)* | 13% (27) | 0% (0) | % (49) | *SW vs E* | < 0.001* |
| “Wild” type | 53% (38) | 46% (96) | 50% (14) | % (148) | *C vs E* | 0.009 |
| TOTAL | 72 | 208 | 28 | 308 | *Full* | < 0.001* |
| *Mutation* | *Change* | *Count* | *Hetero* | *Homo* | *% Total (n = 79)* |  |
| Leu128Ser<br>(L128S) | TTA>TCA | n = 31 | 16/28 | 15/28 | 19% (15/79) |  |
| Tyr139Cys<br>(Y139C) | TAT>TGT | n = 63 | 16/64 | 47/64 | 62% (49/79) |  |

A weighted median of 9% and 82% of the alleles at codons 128 and 139, respectively, were resistant at each capture site (combined x = 50%). All but one sample (78/79) contained at least one copy of a known anticoagulant resistance allele (99%). This fully “wildtype” individual was from London, in Southwestern Ontario. Double resistant individuals (L128S + Y139C, Goulois et al. 2017) were found in all regions except Quebec, as L128S was absent in this region (Fig. 1). No “double homozygous” resistant individuals were found, and no homozygous/heterozygous resistant individuals were detected. When L128S occurred independent of Y139C it was almost always in homozygous form. If the allele was heterozygous, it was always in association with a heterozygous Y139C, except in one case of an L128S heterozygote from Sudbury.

### Allelic & Genotypic Frequencies

L128S was found at eight sites, and in all regions except Quebec, including as far north as Sudbury (Val Caron, Northern Ontario). Y139C was found at every site except Sudbury. There was a significant difference in resistant allele frequencies between regions, except between Central and Eastern (p > 0.05, Table 3). L128S was significantly more common in Southwestern (p < 0.01), as was Y139C in Central (p < 0.001, Tab. 3). Similarly, genotypic frequencies varied significantly between all regions, except between Central and Eastern (Tab. 4). The most common genotype observed was Y139C homozygous in Central Ontario (73%), followed by double heterozygotes and L128S homozygotes in Southwestern Ontario (39% and 44%, respectively). Central Ontario had significantly more Y139C homozygotes than Southwestern (73% vs 11%), and Southwestern had significantly more L128S homozygotes than central (44% vs. 10%, Fisher’s test w/ Bonferronni adjustment p < 0.001, Tab. 4).

**Table 4:** Genotypic counts & frequencies between regions. Genotypic frequencies differed between all regions (except between Central and Eastern). Bonferroni comparisons are denoted by grey shading and asterisks. The only pairwise difference was between the Y139C and L128S homozygous genotypes, in Southwestern and Central Ontario (both p < 0.005). Superscript “W” indicates that allele is wild-type at the locus, bold “R” indicates resistant (e.g. L128S^W/**R**^ / Y139C^W/**R**^ is heterozygous for resistance at both loci).

|  | S. Western | Central | Eastern | TOTAL | Location(s) |
| --- | --- | --- | --- | --- | --- |
| 128<sup>W/R</sup> / 139<sup>W/R</sup> | 39% (7) | 13% (7) | 0% | 18% (14) | Vaughan,<br>Toronto,<br>London |
| 128<sup>R/R</sup> / 139<sup>W/W</sup> | 44% (8)* | 10% (5) | 0% | 17% (13) | Sudbury,<br>Toronto,<br>Brampton,<br>London |
| 128<sup>W/W</sup> / 139<sup>R/R</sup> | 11% (2) | 73%* (38) | 100% (7) | 61% (47) | Toronto,<br>London,<br>Quebec |
| 128<sup>W/W</sup> / 139<sup>W/R</sup> | 0% (0) | 4% (2) | 0% | 3% (2) | Toronto |
| 128<sup>W/W</sup> / 139<sup>W/W</sup> | 6% (1) | 0% | 0% | 1% (1) | London |
| TOTAL | 18 | 52 | 7 | 77 |  |
| *Fisher's p* < 0.001* | *SW vs C* < 0.001* |  | *SW vs E* < 0.001* |  | *C vs E* = 0.83 |

Homozygous resistant individuals of either allele were the most common general genotype, partially due to a lack of wild-type individuals. Haldane’s exact test for Hardy-Weinberg equilibrium demonstrated an excess of L128S heterozygotes in Central Ontario (D = - 3.26, p = 0.002), and an excess of Y139C heterozygotes in the Central region (D = -3.61, p = 0.007), with an excess of both genotypes when regions were combined (Tab. 5).

**Table 5:** Genotypic frequency testing for Hardy-Weinberg equilibrium within and across regions with Haldane’s exact test. Both heterozygous genotypes were more common in Central Ontario, and between population differences were low.

|  | *Y139C* | *L128S* |
| --- | --- | --- |
| Southwestern ON | D = -0.32 | D = -0.65 |
| n = 18 | p = 1 | p = 0.61 |
| Central ON | D = -3.26 | D = -3.61 |
| n = 52 | p = 0.007* | p = 0.002* |
| Ontario-Wide | D = -7.7 | D = -7.81 |
| n = 70 | p < 0.001 * | p < 0.001* |

Linkage disequilibrium was tested within and across both regions to determine if these resistant SNPs were frequently inherited together (Tab. 6). D’ and R^2^ values for the two loci indicate strong linkage disequilibrium and correlation in Southwestern Ontario and Central Ontario, as well as province-wide (all D’= 0.99, p < 0.001), signifying strong historical linkage between L128S and Y139C in both regions (Tab. 6).

**Table 6:**
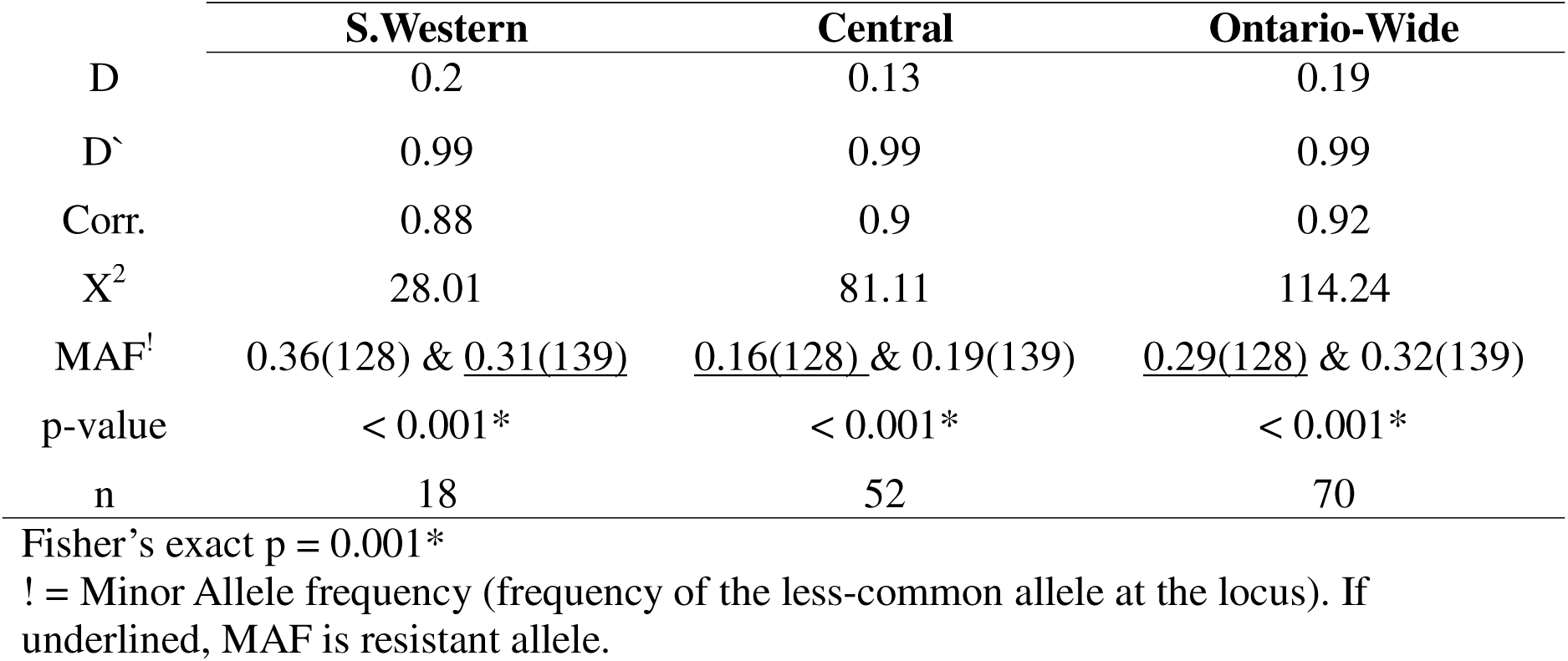
VKORC1 Linkage disequilibrium in house mice across and within regions of Ontario. Across the province, Y139C and L128S are rarely inherited separately, but the degree of LD decreases in Southwestern Ontario, possibly as a function of landscape type (less developed) or population density (lower density).

|  | **S.Western** | **Central** | **Ontario-Wide** |
| --- | --- | --- | --- |
| D | 0.2 | 0.13 | 0.19 |
| D' | 0.99 | 0.99 | 0.99 |
| Corr. | 0.88 | 0.9 | 0.92 |
| X<sup>2</sup> | 28.01 | 81.11 | 114.24 |
| MAF<sup>!</sup> | 0.36(128) & <u>0.31(139)</u> | <u>0.16(128)</u> & 0.19(139) | <u>0.29(128)</u> & 0.32(139) |
| p-value | < 0.001* | < 0.001* | < 0.001* |
| n | 18 | 52 | 70 |

Fisher's exact p = 0.001\*
! = Minor Allele frequency (frequency of the less-common allele at the locus). If underlined, MAF is resistant allele.

### Correlates of Resistance: Linear Regression & Multivariate Analysis

Log-log linear regression analyses were conducted to predict the percentage of individuals that were homozygous resistant at either locus combined, using sites with at least four samples. A total of eight capture sites across Southwestern, Central Ontario and Quebec were used for this purpose (Tab. 7). These were initially included as single predictors only, in four separate linear models, to prevent multicollinearity. The population density in the census tract of the capture site was the only predictor of homozygous resistance. The percentage of developed land (residential + industrial + commercial) was trending toward significance (Tab. 7). Median income and post-secondary education rate were also insignificant. Cook’s distance showed that, in the *Population density* model, the site *London* was moderately influential (∼0.65), and in the *Developed land* model, *London* was highly influential (∼0.8). This site had the second lowest population density and second lowest percentage of developed land, next to that of *Wyoming,* Ontario (Fig. 2).

**Figure 2:**
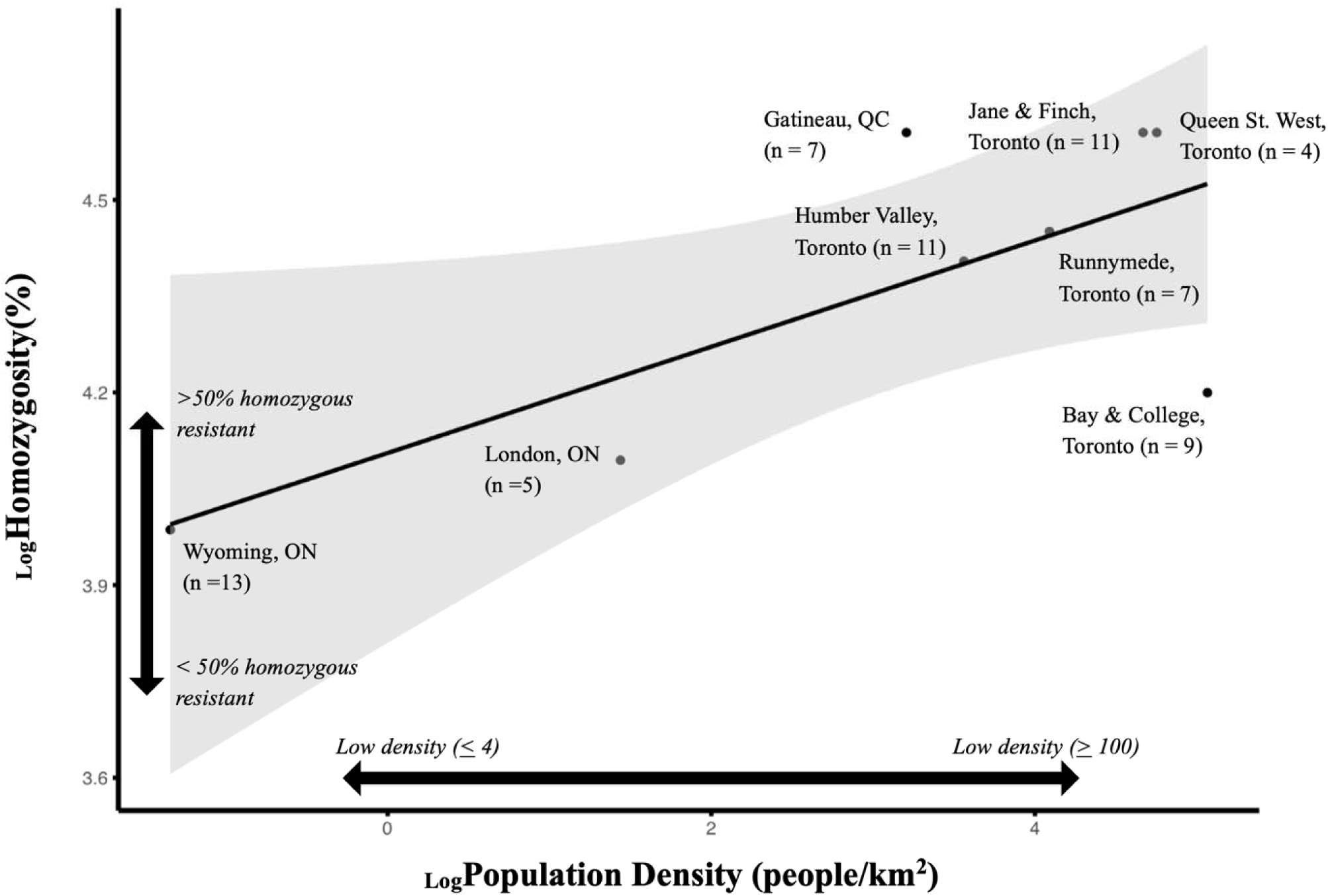
Log-log linear regression; correlation between L128S and Y139C (log)homozygosity as a function of (log)population density. Only sites with at least four observations were used (n = 8 sites, 67 samples). Shaded region is 95% confidence interval. P = 0.04, R^2^ = 0.53.

**Table 7:** Log-log linear model selection for _Log_ homozygous resistance. Linear regression was used to predict the percentage of individuals that were homozygous resistant at either codon 128 or 139. The strongest and best-fitted single predictor was population density in the census tract of the capture site; n = 8 sites, 67 samples.

|  | Coef. | p-value<br>(slope) | Pearson's R | R<sup>2</sup> | AIC |
| --- | --- | --- | --- | --- | --- |
| $\text{Log}\%$ Post secondary Education<br><u>Census tract</u> | -1 | 0.28 | -0.44 | 0.19 | 3.54 |
| $\text{Log}$ Med. Income (\$/CAD)<br><u>Census tract</u> | -0.2 | 0.59 | -0.23 | 0.05 | 4.82 |
| $\text{Log}$ Population density (people/km<sup>2</sup>)<br><u>Census tract</u> | 0.1 | 0.04* | 0.73 | 0.53 | 1.06 |
| $\text{Log}\%$ Developed land<br><u>ArcGIS Online</u> | 0.98 | 0.089 | 0.64 | 0.41 | 1.63 |
| **PC1** of all predictors | 6.72 | 0.19 | 0.52 | 0.27 | 72.11 |

A Principal component analysis of all four predictors produced a PC1 explaining 53% of variance, which, when correlated with the percentage of homozygous resistant individuals at a site, was insignificant (Tab. 7). PC loadings demonstrated that, as the percentage of developed land decreased at a capture site concomitantly with population density within the census tract, median income and post-secondary education rate increased (Tab. 7), corresponding with a decrease in the percentage of resistant homozygotes in the population (Fig. 2). Loadings for PC1 were 0.58 (developed land), 0.55 (pop. density), -0.58 (median income) and -0.19 (education rate).

## Discussion

We sought to determine if VKORC1 rodenticide resistance SNPs were present in Ontario house mice and examine their regional allelic and genotypic variability, as well as correlation with socioeconomic and land usage factors. We recorded substantial frequencies of known resistance SNPs in wild house mice caught across the province of Ontario, distinct patterns of allelic and genotypic occurrence between regions, and an increase in homozygous resistant mice in high population density census tracts. Genetic rodenticide resistance in house mice has not, up until now, been examined in Canada, much less with respect to regional, socioeconomic and environmental variation.

The two SNPs we found (L128S and Y139C) have been thoroughly documented worldwide (USA, England, France, Italy, Netherlands, Germany, Russia, North Africa, Middle East, Hong Kong, etc., McGee et al. 2020; Díaz and Kohn 2021b; Maltsev et al. 2022). L128S (Leu128Ser, “Cambridge Cream” resistance) was first documented in Cambridge, England (Wallace and MacSwiney 1976), whereas Y139C (Tyr139Cys, “Berkshire” or “Reading” resistance) was first documented in house mice from Reading, England (Pelz et al. 2005; Goulois et al. 2017). Both are known to cause extreme practical resistance to first-generation ARs. Y139C is also known to cause bromadiolone (SGAR) resistance, increasing lethal dosage by a factor of 5.6-21 (Baxter et al. 2022) based on blood-clotting response tests (BCR, Martin et al. 1979), *in-vivo* studies and field trials (Pelz et al. 2012). Practically resistant mice are those which require at least ten times the dosage of AR active ingredient for lethality, which poses serious issues for field efficacy (Baxter et al. 2022).

Anticoagulant resistance (*in-vivo*) was last examined in Toronto, Ontario in 1982 (Siddiqi and Blaine 1982), and only two other times in Canada (Cameron 1977 unpublished data, Cronin 1979). The earliest attempt was in 1977 by PCO Services (Cameron 1977 unpublished data). These mice were collected from regions of Toronto comparable to some of those from which we obtained samples (Siddiqi 2023, personal communication). Given that anticoagulant usage has not stopped since it began in Ontario, we assume that resistance has likely only increased overall. In our sample of 49 mice from Toronto, as well as three from the Greater Toronto Area (Vaughan and Brampton), we found 100% resistance, in that all samples had at least one resistant allele at either locus. Most of the genotypes we found are known to confer heavy practical resistance even to the most popular second-generation AR in the country; bromadiolone (Elliott et al. 2014; Baxter et al. 2022). In 1980s Toronto (Siddiqi and Blaine 1982), 71.43% of house mice were resistant to warfarin, with 96% of these also being cross-resistant to the first-generation AR chlorophacinone. However, the fact that their LD50 for bromadiolone (1.75 mg/kg, Siddiqi and Blaine 1982) was quite similar to that discovered recently for susceptible house mice in Britain (1.96 mg/kg, Baxter et al. 2022) but their warfarin resistance (LD50 374 mg/kg, Siddiqi and Blaine 1982) corresponds more so with that of homozygous resistant Y139C (412.16 mg/kg, (Baxter et al. 2022) is puzzling, given that heavy resistance to first-generation ARs should provide moderate cross-resistance to bromadiolone (Mooney et al. 2018). However, these LD50s were calculated using feeding tests and BCR, respectively, which may provide different results, and BCR is possibly less reliable for SGAR toxicity (Pelz et al. 2005). Regardless, Toronto mice may have formerly been achieving resistance by another mechanism, such as gut flora vitamin K synthesis or detoxification (Markussen et al. 2008; Poché and Poché 2012), and may have only gained resistance SNPs after the 1980s. Assuming that this resistance was due to these same VKORC1 SNPs, the resistant mice would likely have possessed some genotype of L128S (Blažić et al. 2018), or perhaps heterozygous Y139C. A delayed time to death after bromadiolone exposure (17-21 days, Siddiqi and Blaine 1982) was noted, indicating the possibility of some acquired tolerance in these mice, as opposed to true “resistance” (Frantz and Madigan 1998).

Resistant house mice are far more common in the UK (the “birthplace” of resistance) than wild-type individuals, as is the case for other regions of Europe (Ruiz-López et al. 2022), indicating that our high prevalence rate is not unusual. Indeed, resistance alleles are commonly present in the majority of samples taken across large geographic areas (58% in Lebanon, Rached et al. 2022, 78% in Ireland, Mooney et al. 2018, 70% in France, Goulois et al. 2017, 90% in Michigan, Díaz and Kohn 2021), ours being similar to those sampled in the American Northeast and Midwest (Díaz and Kohn 2021). Resistance is known to occur and become common after five to ten years of “sustained” anticoagulant usage acting as a selection pressure, with homozygotes proliferating under periods of intensive AR usage (Kohn et al. 2000). Given this timeline and the prevalence of homozygous resistant individuals in our samples, it appears evident that house mice across the province have consistently been exposed to ARs for numerous decades, which corresponds with Canadian practices of “permanent baiting” (Hindmarch et al. 2018) that often occur on urban commercial properties (Government of Ontario 2017). However, given the worldwide increase in urbanization since the mid-20th century (Tripodi et al. 2024), and the correlation we have detected between homozygous resistance and population density, resistance may now be spreading at an even greater rate in Ontario. It may also be true that samples donated by PMPs are simply more likely to be resistant. Considering PMP’s widespread usage of ARs which select for resistance, in combination with the traps which were used to collect these samples, this high prevalence may simply be the result of a sampling bias (Byers et al. 2019).

AR resistance was noted in US populations of house mice and Norway rats beginning in the 1970s (Brooks and Bowerman 1973) and has since been documented throughout the country, and in three states that border Ontario (Minnesota, Michigan and New York), with an eastward increase in genotypic prevalence (Díaz and Kohn 2021). Of the samples taken in the nearby USA, New York State resistant house mice largely possessed Y139C, with only one independent L128S individual (n = 86). Moving further west, house mouse samples from Michigan (closest to Southwestern Ontario) were resistant (90%) largely as a result of L128S. Those from Illinois also lacked Y139C, but possessed L128S (15.4%). Resistance prevalence in Minnesota was also noticeably lower than that of Michigan (42.1%), and was largely due to L128S. Knowing this, and considering that no resistance alleles were detected in those states closest to our samples from Gatineau, Quebec (Vermont & New Hampshire, Díaz and Kohn 2021), this suggests two separate branches of wild house mouse geneflow between Canada and the United States for each of the two resistant loci, or at least, a similar pattern of selection pressure leading to similar genotypic frequencies: 1. Midwestern USA/Southwestern Ontario, for L128S, and 2. Northeastern USA/South Central Ontario, for Y139C.

A previous study found that urban populations of rodents tend to have greater allele fixation than rural populations (Harris and Munshi South 2017). We observed that our two sites Southwestern Ontario were far more undeveloped than those from Central Ontario, and homozygous resistance was less common in Southwestern. Little information on the relatedness of house mouse populations on a geographic scale has been conducted, but resistance in rats is said to be significantly related to location in European populations (Desvars-Larrive et al. 2017), as well as land usage (Marquez et al. 2019). It is unknown if the same is true for house mice. Considering that L128S and Y139C were in strong linkage disequilibrium in our samples, and that L128S was absent from Quebec further suggests two pathways of spread or regional selective pressures for these strains, and dominance of the Y139C allele, which has been found previously in other regions (Aivelo et al. 2025).

The SNPs we found may have spilled over from individuals from European populations (Díaz and Kohn 2021), but may also have arisen independently (Rost et al. 2009, Díaz and Kohn 2021). L128S and Y139C are the two most widespread resistance SNPs in North American and European house mice, and are known to frequently occur together. In rats from Germany, resistance has been shown to be strongly correlated with neutral microsatellite loci (Kohn et al. 2000), and rats in England are experiencing some degree of structure associated with microsatellites and VKORC1 mutations (Haniza et al. 2015). This suggests that intensive rodenticide usage may lead to a selective sweep as a result of extreme environmental disfavour of the VKORC1 “wild-type” and its associated genetic diversity in rats, and possibly for house mice also.

We found more homozygous resistant individuals (for L128S and Y139C combined) as a function of greater population density, which is consistent with the results of previous studies (Marquez et al. 2019). The proportion of resistant individuals or resistant alleles were not modelled due to lack of variation. Few studies have attempted to directly correlate land usage (as a possible proxy for rodenticide usage, Buckley et al. 2024) or other predictors with the prevalence of resistance. However, population density (Geduhn et al. 2015; López-Perea et al. 2015; Buckley et al. 2024) and urbanization (Burke et al. 2021) have been associated with a greater probability of non-target exposure, and with resistance (Marquez et al. 2019), just as lower income communities are known to be disproportionately afflicted by rodent infestations (Feng and Himsworth 2014). Our PCA loadings demonstrate multicollinearity between population density and developed land, as well as an inverse correlation with education rate and median income. Highly dense, urban and low-income communities may represent a double-edged sword for rodent control, having greater commensal populations as well as greater homozygous resistance. Although homozygosity decreases susceptibility to ARs by a large margin, it is also said to drastically increase K requirements by reducing the VKORC1 receptor’s affinity for ARs and vitamin K simultaneously (Debaux et al. 2019), thus shortening the animal’s lifespan (Smith et al. 1991) and possibly decreasing fecundity (Heiberg et al. 2006). Therefore, a moratorium on SGARs could allow for a reduction in resistance, after which populations could be quickly extirpated by the most potent SGARs (Aivelo et al. 2025). Some previous studies do suggest that resistance decreases quickly after baiting cessation (Quy et al. 1995, Cowan et al. 2017), but others in the past have contradicted this (Rowe and Redfern 1965, Partridge 1979).

L128S is considered to provide less resistance than Y139C, but may be more prevalent in Southwestern Ontario and the American Midwest because of different AR usage patterns, not as a result of gene flow (Heiberg et al. 2006), which could be related to different land usage composition. However, resistance alleles may still become fixed and remain long after the cessation of AR usage (Heiberg et al. 2006). Regardless, if chemical control of house mice continues as is, the two strongest second-generation ARs in Ontario (difethialone and brodifacoum) will be necessary for control, since there is supposedly little practical resistance to these chemicals (Blažić et al. 2018). If bromadiolone remains the main AR used, resistance may be exacerbated by a management feedback loop, causing pest controllers to use greater quantities of the same chemicals selecting for resistance (Brakes and Smith 2005).

Although AR usage will increase the frequency of resistance (Pelz et al. 2012), resistance may still occur in some populations of wild rodents and commensals not exposed to ARs (Díaz et al. 2010, Sran et al. 2023) especially house mice (Baxter et al. 2022). Regardless, using ARs in an uncoordinated and piecemeal fashion, or applications by untrained individuals (Goulois et al. 2017) before coordinated management is implemented, may select for resistant alleles, making large-scale AR control efforts difficult (Gallozzi et al. 2024). As such, low-income communities (Himsworth et al. 2014), and highly urbanized (Tripodi et al. 2024) high-density areas (De Cock et al. 2024) may experience greater challenges with rodent control, likely resulting in part from resistance. This could be the result of nearby permanent baiting which occurs in commercial areas, often adjacent to low-income residences (Hindmarch et al. 2018), contributing to resistance proliferation, or greater usage of rodenticides in lower income regions (Murray et al. 2025). Although permanent baiting is a longstanding practice (Rowe and Swinney 1988), and had excellent control effects early on (Lund, M. 1972), it is now likely a major cause of resistance and non-target exposure (Hindmarch et al. 2018). Our PCA, which showed decreased income and education rate concomitant with increased population density and developed land, reflects this pattern well, however, only developed land was significantly correlated with homozygous resistance.

Admittedly, our Sanger sequencing success (66%) may have been influenced by the age of the samples, since some mice had been dead for weeks before collection by PMPs. The quality of the freezers used for storage at PMP offices may have also had an effect. However, qPCR success was 100% (79/79), indicating that this method is far superior. The qPCR program also provided definitive genotype calls for the samples along with the level of confidence. Therefore, we recommend qPCR assays for known resistance codons as an efficient way of monitoring VKORC1 resistance SNPs in house mouse populations, especially when samples are collected from PMPs (Krijger et al. 2023).

## Conclusion and Future Directions

Based on our findings, it is evident that pest management practices have made great impacts on house mouse populations in Ontario, and other traits may be linked to AR resistance which we did not detect or investigate. It seems likely that FGARs will be wholly insufficient for house mouse control throughout much of Ontario, and possibly Quebec. Bromadiolone may be effective in regions of lower population density, or areas with less intensive control efforts, due to the lower prevalence of homozygous resistance. Therefore, the strongest available SGARs (difethialone, brodifacoum) or non-anticoagulants, may be necessary for chemical control of house mice in much of Ontario, even where only heterozygous resistance is prevalent. Future research should seek to identify the origins of these resistance SNPs through microsatellite analysis (Kohn et al. 2000; Haniza et al. 2015), which may suggest gene flow between provinces or countries as a result of stowaway house mice travelling between regions. This may help to establish international management campaigns for this highly invasive and destructive species.

## Acknowledgements

The authors would like to thank Dr. Claire Jardine and Dr. Jason Munshi-South for their initial reviews of the thesis from which this manuscript was derived. Pest management companies have chosen to remain anonymous, but we would like to thank their management teams and technicians for donation of samples.

## Author Contributions

L. Richardson, A. Schulte-Hostedde and C. Cullingham contributed to the study conception and design. Material preparation, data collection and analysis were performed by L. Richardson. All authors read and approved the final manuscript. DNA amplification and qPCR was performed by C. Cullingham, M. Balsara and C. Larose. The first draft of the manuscript was written by L. Richardson, and all authors commented on and added to/edited previous versions of the manuscript.

